# Subchronic Toxicity Study of Nitric Oxide Nanobubbles Injection in Sprague-Dawley Rats

**DOI:** 10.64898/2026.01.10.698772

**Authors:** Dody Novrial, Nor Sri Inayati, Nur Signa Aini Gumilas, Dhadhang Wahyu Kurniawan, Sutiman Bambang Sumitro

## Abstract

Following the success of hydrogen nanobubble technology, development has expanded to include similar products with other gases, such as nitric oxide (NO). In previous studies, NO donors have been frequently used for both therapeutic and diagnostic purposes. However, the use of NO in nanoformulations has received little attention thus far. Thus, 90 days of subchronic toxicity testing in Sprague-Dawley rats were used in this study to assess potential adverse effects of intravenous administration of NO nanobubbles (NONB). Hematological factors, serum biochemistry, and histology of the liver, kidney, heart, lung, and spleen were investigated. The findings of the intravenous NONB injection trial in rats, using graded doses of 0.01 mL, 0.04 mL, and 0.06 mL, demonstrated no deaths during the 90-day treatment period. Biochemical parameters for liver function, lipid profile, kidney function, and serum electrolyte levels remain within acceptable limits, with slight variations in sodium and potassium levels. This condition was also supported by histological findings in the liver, kidney, and spleen, indicating mild to moderate damage, while the heart and lungs were normal. Thus, the intravenous administration of NONB at a dose of up to 0.06 mL in this subchronic toxicity test remains safe. Further studies with adjustments to the test formulation are highly encouraged until the clinical trial stage in humans.

## Introduction

Nanobubble technology has been extensively developed for medical and pharmaceutical applications. Existing research demonstrates promising results, especially regarding its stability and permeability(1,2). These bubbles, due to their small size, can pass through cell membranes more easily, making them suitable for targeted therapies(3). Hydrogen nanobubble technology is an example of cutting-edge innovation that harnesses hydrogen at the nanoscale. These tiny bubbles possess unique properties that make them highly effective across a range of applications, from energy storage to medical treatments(4). This technology enables the distribution of hydrogen to multiple systems with greater stability and efficiency by encapsulating the gas in tiny bubbles(5).

Building on the success of hydrogen nanobubble technology, development has expanded to similar products using other gases, including Nitric Oxide (NO). NO is a simple gas composed of two atomic elements: oxygen (O) and nitrogen (N). It is colorless and odorless, dissolving rapidly in both organic and aqueous solvents. The enzyme nitric oxide synthase (NOS) catalyzes the production of NO in various organisms, including bacteria, fungi, plants, and animals, as part of their cellular metabolic processes(6).

NO donors were commonly used in earlier research for both medicinal and diagnostic purposes. Microbubbles containing NO have been shown to enhance vasodilation, decrease platelet aggregation, and lower the likelihood of inflammatory cell infiltration and thrombus formation. Myocardial infarction and stroke management and prevention greatly benefit from this ability(7–9). However, the application of NO in nanoformulations has not been extensively studied to date. Our previous research, involving the intravenous infusion of a solution containing NO, Mg, and hydrogen nanobubbles, confirmed a safe level with no significant side effects. However, an increase in AST, ALT, and urea levels was observed in the treatment group, which correlated linearly with increasing NO doses and decreasing hydrogen nanobubble doses in the test preparation(10). These results indicate that NO in the injection solution is an important factor in increasing AST, ALT, and urea levels. Thus, this work uses subchronic toxicity testing in Sprague-Dawley rats to evaluate the potential health risks associated with the injection of NO nanobubbles (NONB).

## Materials and Methods

### Injection material

The injection material was obtained from the Indonesian Molecular Innovation (IMI) Foundation as an infusion fluid containing NONB at a concentration of 26 million bubbles/mL. The NONB solution was prepared in reverse osmosis. A buffer containing 1% ascorbic acid and 1% sodium bicarbonate was used to maintain pH stability. All materials were prepared under laboratory protocols, and all preparations were conducted aseptically to minimize contamination(11).

### Animal material

This study was conducted on male and female Sprague-Dawley rats, aged 8-12 weeks and weighing 150-250 g at the start of the experiment. Females (nulliparous and not pregnant) and males were separated in cages during the study. The animals were acclimated for one week before the start of the experiment. The animals had free access to tap water for drinking and to standard laboratory feed. The rats were housed in groups of 5 per cage at room temperature (25 ± 3 °C), with sufficient ventilation and a natural light cycle (12 hours/12 hours). All procedures and protocols involving animals and their care were conducted in accordance with the guidelines of the Research Laboratory of the Faculty of Medicine, Universitas Jenderal Soedirman, and are approved by the Ethics Committee of the Faculty of Medicine, Universitas Jenderal Soedirman (Reg. No. 065/KEPK/PE/VIII/2024).

### Evaluation of subchronic toxicity

The subchronic toxicity of NONB injection was evaluated according to the modified OECD guideline 408(12). Six groups of 14 animals each (7 males and 7 females) were used for each dose. Group 1 served as a healthy control, and groups 2, 3, and 4 received NONB intravenous injections into the tail vein (0.01mL, 0.04 mL, and 0.06 mL, respectively). Group 5 was a healthy satellite control. Group 6 was the satellite group that received an intravenous injection of 0.06 mL NONB into the tail vein. Those two groups were observed for an additional four weeks after the other groups. During treatment (90 days), the animals’ body weights were assessed weekly, and signs of toxicity were carefully examined.

### Blood and organ collection

Blood samples were collected once at the end of the experiment, just before termination. The animals’ blood samples were collected after they had fasted overnight (10 hours). It was extracted from the retro-orbital vein and stored in non-EDTA tubes for biochemical analysis and in EDTA tubes for haematological studies. Then, the rats were terminated immediately. After dissecting the rats, organs such as the heart, liver, lungs, kidneys, and spleen were removed, cleaned in saline (0.9%), blotted on absorbent paper, and weighed to determine relative organ weight. The organs were then fixed in buffered formalin for histological examination.

### Blood analysis

White blood cells (WBC), red blood cells (RBC), lymphocytes (LYM), monocytes (MON), neutrophils (NEUT), eosinophils (EO), basophils (BASO), hemoglobin (HB), hematocrit (HCT), and platelet count (PLT) were all measured. Meanwhile, serum biochemical analysis was performed, including glucose, total cholesterol, triglycerides, ALT, AST, urea, creatinine, sodium, and potassium.

### Histological analysis

The sample organs were dried and fixed in paraffin for microscopic inspection using standard pathology laboratory techniques. Histological investigation was performed using paraffin sections stained with hematoxylin and eosin (H&E). Histological scoring was performed in 10 fields of view per organ sample at 100x magnification. Liver histology was assessed according to the following criteria: normal liver (score 1); focal hydropic/ fatty degeneration/ necrosis (score 2); multifocal hydropic/ fatty degeneration/ necrosis (score 3); diffuse hydropic/ fatty degeneration/ necrosis throughout the entire field of view (score 4). Kidney histology was assessed according to the following criteria: normal kidney (score 1); lesions such as nuclear necrosis, tubular degeneration, and proximal tubular dilation were found in some fields of view (score 2); lesions such as nuclear necrosis, tubular degeneration, and proximal tubular dilation were found in every field of view (score 3). Heart histology is assessed according to the following criteria: normal heart (score 1); mild lesions (inflammation/fibrosis/vacuolization < 3 foci) (score 2); moderate lesions (inflammation/fibrosis/vacuolization > 3 foci) (score 3); severe lesions (diffuse inflammation/fibrosis/vacuolization) (score 4). Lung histology was assessed based on the following criteria: normal lung (score 1); alveolar wall thickening without lung structure damage (score 2); alveolar wall thickening/fibrosis with lung architecture damage (score 3); extensive fibrosis with a honeycomb appearance (score 4); total lung damage (score 5). Spleen histology was evaluated according to the following criteria: normal spleen (score 1); focal hemorrhage/necrosis (score 2); multifocal hemorrhage/necrosis (score 3); diffuse hemorrhage/necrosis (score 4).

### Statistical analysis

Continuous data are presented as mean ± SD, while categorical data are presented as numbers (percentages). The ANOVA and Kruskal-Wallis test, followed by the appropriate post hoc test, were used in statistical analysis. A p-value < 0.05 indicates a significant difference in the variable’s mean.

## Results

### Effects of NONB on the morbidity and mortality of rats

Observations made during the investigation indicate that no experimental animals perished. None of the test animals had convulsions, tremors, or diarrhea. No discernible behavioral alterations were observed during physical examination; the experimental animals’ activity levels did not decline. The condition of the fur of the experimental animals did not alter, according to the observation (Table 1).

**Table 1.**
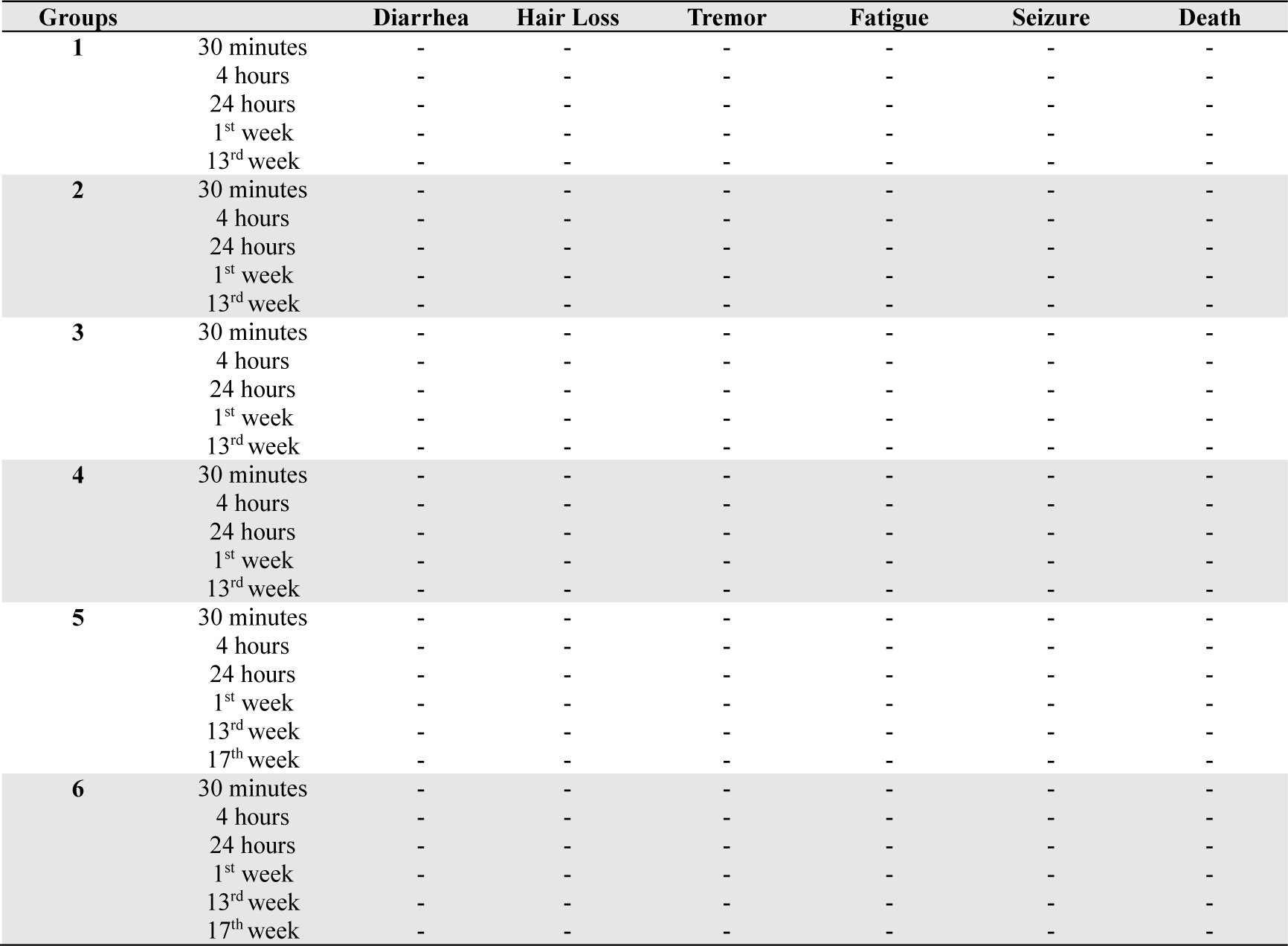
Effects of NONB on the morbidity and mortality of rats.

### Effects of NONB on the relative organ weight and body weight of rats

The effects of NONB on the relative weights of some organs are summarized in Table 2, and the evolution of body weights of rats is displayed in Figure 1. The average relative organ weights of the rats were not significantly affected by the test substance, as the organ weights in the treatment group did not differ substantially from the normal organ weight range. Throughout the experiment, the rats consistently gained weight. The average weight of the rats at the beginning of the study was around 180 grams, while by the end, it had risen to nearly 290 grams.

**Figure 1.**
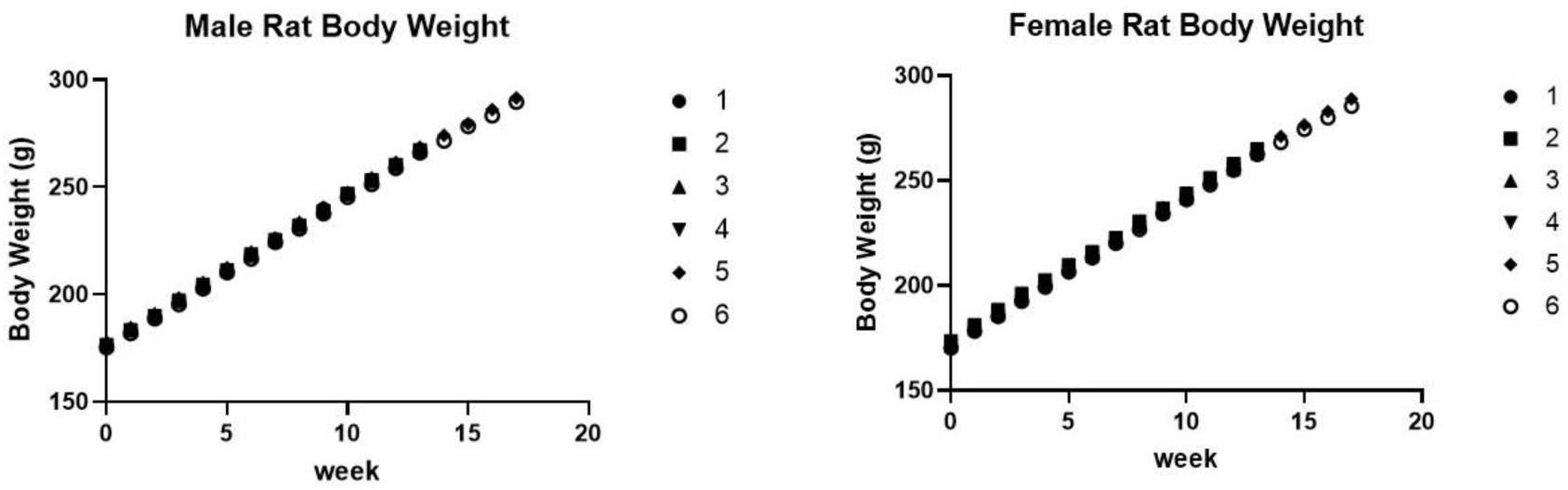
Evolution of the rat’s body weight

**Table 2.**
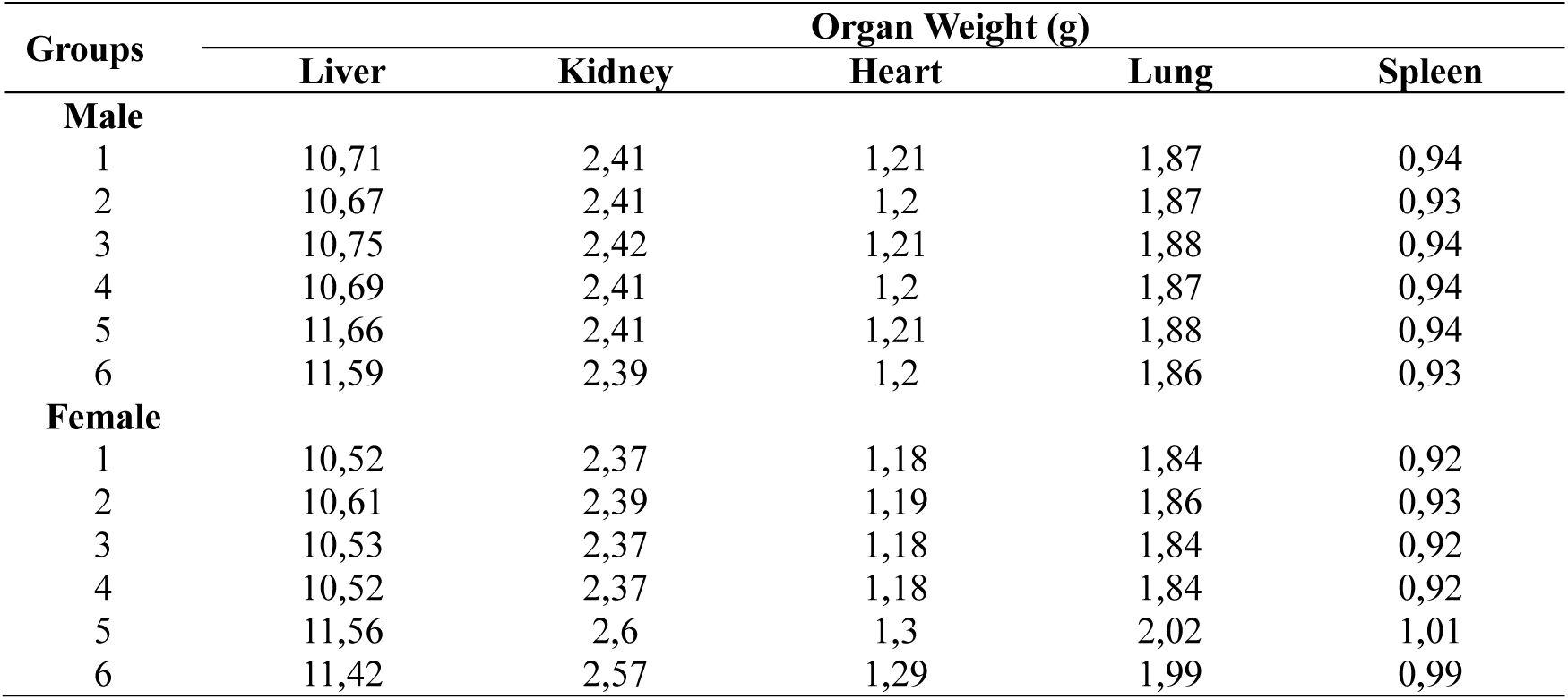
The relative organ weight of rats.

| Groups | Organ Weight (g) |  |  |  |  |
| --- | --- | --- | --- | --- | --- |
|  | Liver | Kidney | Heart | Lung | Spleen |
| **Male** |  |  |  |  |  |
| 1 | 10,71 | 2,41 | 1,21 | 1,87 | 0,94 |
| 2 | 10,67 | 2,41 | 1,2 | 1,87 | 0,93 |
| 3 | 10,75 | 2,42 | 1,21 | 1,88 | 0,94 |
| 4 | 10,69 | 2,41 | 1,2 | 1,87 | 0,94 |
| 5 | 11,66 | 2,41 | 1,21 | 1,88 | 0,94 |
| 6 | 11,59 | 2,39 | 1,2 | 1,86 | 0,93 |
| **Female** |  |  |  |  |  |
| 1 | 10,52 | 2,37 | 1,18 | 1,84 | 0,92 |
| 2 | 10,61 | 2,39 | 1,19 | 1,86 | 0,93 |
| 3 | 10,53 | 2,37 | 1,18 | 1,84 | 0,92 |
| 4 | 10,52 | 2,37 | 1,18 | 1,84 | 0,92 |
| 5 | 11,56 | 2,6 | 1,3 | 2,02 | 1,01 |
| 6 | 11,42 | 2,57 | 1,29 | 1,99 | 0,99 |

### Effects of NONB on the hematological parameters

The effects of NONB on some of the hematological parameters are recorded in Table 3 (male) and Table 4 (female). Tables 3 and 4 show that NONB administration did not affect blood parameters in male and female rats.

**Table 3.** Hematological parameters of male rats.

| Parameters | Groups |  |  |  |  |  | *p* |
| --- | --- | --- | --- | --- | --- | --- | --- |
|  | 1 | 2 | 3 | 4 | 5 | 6 |  |
| Number of Erythrocytes (10<sup>6</sup>/mm<sup>3</sup>) | 8.39 ± 0.37 | 8.50 ± 0.46 | 8.04 ± 0.30 | 8.00 ± 0.18 | 8.21 ± 0.31 | 8.66 ± 0.45 | 0.74 |
| Hematocrit (%) | 41.71 ± 1.48 | 43.16 ± 2.03 | 47.49 ± 1.98 | 41.77 ± 2.19 | 43.12 ± 1.38 | 43.93 ± 2.62 | 0.36 |
| Hemoglobin (g/dl) | 15.55 ± 0.45 | 15.18 ± 0.37 | 15.68 ± 0.28 | 15.71 ± 0.30 | 14.87 ± 0.33 | 15.57 ± 0.32 | 0.44 |
| Number of leukocytes (10<sup>3</sup>/mm<sup>3</sup>) | 11.41 ± 1.91 | 14.51 ± 1.42 | 12.04 ± 1.69 | 12.14 ± 1.77 | 12.38 ± 1.57 | 13.25 ± 1.74 | 0.82 |
| MCH (pg) | 17.18 ± 0.38 | 18.01 ± 0.59 | 18.27 ± 0.35 | 17.73 ± 0.46 | 17.38 ± 0.47 | 17.98 ± 0.52 | 0.57 |
| MCHC (g/dl) | 34.72 ± 0.97 | 34.66 ± 0.89 | 35.48 ± 0.72 | 35.03 ± 1.04 | 32.81 ± 0.85 | 34.58 ± 0.64 | 0.52 |
| MCV (fl) | 54.49 ± 1.09 | 50.84 ± 1.87 | 54.73 ± 0.99 | 50.11 ± 1.71 | 55.31 ± 2.45 | 54.73 ± 1.91 | 0.22 |
| Methemoglobin (% Hb) | 0.87 ± 0.98 | 0.79 ± 0.11 | 0.89 ± 0.76 | 0.99 ± 0.06 | 0.96 ± 0.63 | 0.68 ± 0.74 | 0.12 |
| Number of Platelets (10<sup>3</sup>/mm<sup>3</sup>) | 941.50 ± 50.94 | 955.83 ± 46.71 | 1042.67 ± 50.61 | 794.50 ± 137.51 | 998.00 ± 56.74 | 970.00 ± 36.57 | 0.33 |

**Table 4.** Hematological parameters of female rats.

| Parameters | Groups |  |  |  |  |  | *p* |
| --- | --- | --- | --- | --- | --- | --- | --- |
|  | 1 | 2 | 3 | 4 | 5 | 6 |  |
| Number of Erythrocytes (10<sup>6</sup>/mm<sup>3</sup>) | 7.81 ± 0.23 | 7.63 ± 0.34 | 7.39 ± 0.27 | 8.29 ± 0.12 | 7.65 ± 0.22 | 7.48 ± 1.60 | 0.132 |
| Hematocrit (%) | 45.33 ± 2.03 | 44.85 ± 1.98 | 44.58 ± 1.51 | 44.57 ± 1.36 | 45.28 ± 1.70 | 43.35 ± 2.15 | 0.975 |
| Hemoglobin (g/dl) | 15.34 ± 0.35 | 15.40 ± 0.27 | 15.44 ± 0.41 | 15.88 ± 0.30 | 15.16 ± 0.23 | 15.31 ± 0.37 | 0.731 |
| Number of leukocytes (10<sup>3</sup>/mm<sup>3</sup>) | 9.36 ± 1.05 | 9.47 ± 1.18 | 10.72 ± 1.25 | 10.20 ± 0.95 | 9.74 ± 1.32 | 9.24 ± 1.03 | 0.937 |
| MCH (pg) | 55.92 ± 1.20 | 54.38 ± 1.14 | 55.67 ± 0.54 | 55.03 ± 1.06 | 55.88 ± 0.45 | 54.3.0 ± 1.31 | 0.750 |
| MCHC (g/dl) | 17.98 ± 0.48 | 17.99 ± 0.55 | 17.48 ± 0.46 | 17.95 ± 0.57 | 17.91 ± 0.52 | 17.78 ± 0.39 | 0.976 |
| MCV (fl) | 34.43 ± 0.99 | 34.65 ± 0.98 | 33.64 ± 0.85 | 34.09 ± 0.98 | 33.82 ± 0.99 | 35.15 ± 0.86 | 0.876 |
| Methemoglobin (% Hb) | 0.91 ± 0.94 | 0.67 ± 0.12 | 0.60 ± 0.03 | 0.59 ± 0.72 | 0.72 ± 0.12 | 0.86 ± 0.13 | 0.140 |
| Number of Platelets (10<sup>3</sup>/mm<sup>3</sup>) | 1083.67 ± 42.61 | 1032.67 ± 59.14 | 994.50 ± 35.45 | 981.83 ± 33.80 | 1042.33 ± 44.89 | 977.50 ± 51.96 | 0.537 |

### Effects of NONB on serum biochemical parameters

The effects of NONB on some of the serum biochemical parameters are summarized in Table 5 (male) and Table 6 (female). Blood glucose levels, total cholesterol, triglycerides, AST, ALT, urea, creatinine, and blood potassium levels in both male and female rats rose as the test preparation dosage increased. Nevertheless, blood sodium levels decreased, and this decrease was inversely correlated with the test preparation’s dosage increase. In both male and female rat groups, the bivariate analysis revealed significant differences across all parameters except urea levels. Post-hoc analyses for glucose, total cholesterol, triglycerides, SGOT, SGPT, creatinine, sodium, and potassium are presented in Table 7.

**Table 5.**
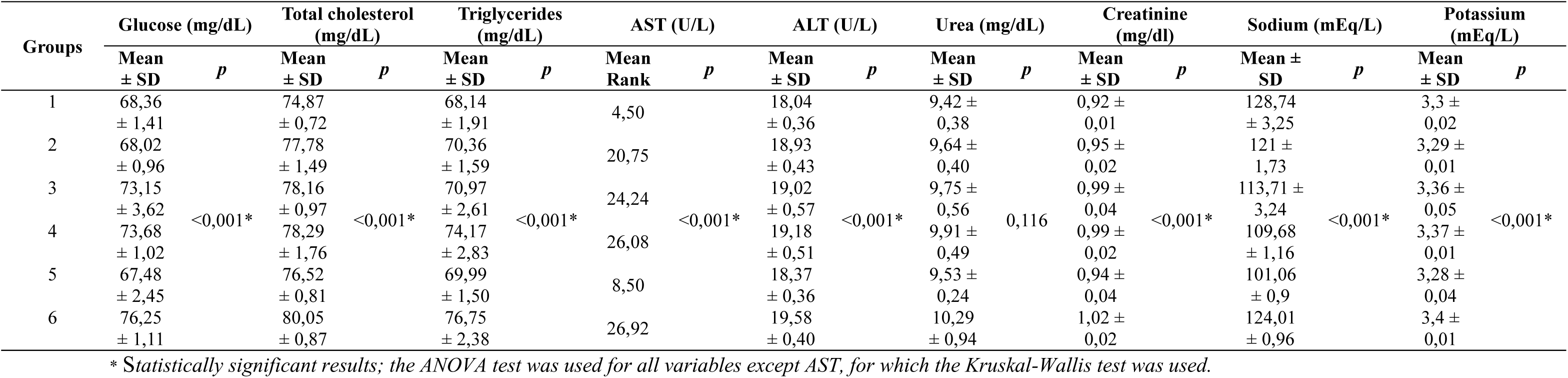
Serum biochemical parameters of male rats.

| Groups | Glucose (mg/dL) |  | Total cholesterol (mg/dL) |  | Triglycerides (mg/dL) |  | AST (U/L) |  | ALT (U/L) |  | Urea (mg/dL) |  | Creatinine (mg/dl) |  | Sodium (mEq/L) |  | Potassium (mEq/L) |  |
| --- | --- | --- | --- | --- | --- | --- | --- | --- | --- | --- | --- | --- | --- | --- | --- | --- | --- | --- |
|  | Mean<br>± SD | *p* | Mean<br>± SD | *p* | Mean<br>± SD | *p* | Mean<br>Rank | *p* | Mean<br>± SD | *p* | Mean<br>± SD | *p* | Mean<br>± SD | *p* | Mean ±<br>SD | *p* | Mean<br>± SD | *p* |
| 1 | 68,36<br>± 1,41 |  | 74,87<br>± 0,72 |  | 68,14<br>± 1,91 |  | 4,50 |  | 18,04<br>± 0,36 |  | 9,42 ±<br>0,38 |  | 0,92 ±<br>0,01 |  | 128,74<br>± 3,25 |  | 3,3 ±<br>0,02 |  |
| 2 | 68,02<br>± 0,96 |  | 77,78<br>± 1,49 |  | 70,36<br>± 1,59 |  | 20,75 |  | 18,93<br>± 0,43 |  | 9,64 ±<br>0,40 |  | 0,95 ±<br>0,02 |  | 121 ±<br>1,73 |  | 3,29 ±<br>0,01 |  |
| 3 | 73,15<br>± 3,62 | <0,001* | 78,16<br>± 0,97 | <0,001* | 70,97<br>± 2,61 | <0,001* | 24,24 | <0,001* | 19,02<br>± 0,57 | <0,001* | 9,75 ±<br>0,56 | 0,116 | 0,99 ±<br>0,04 | <0,001* | 113,71 ±<br>3,24 | <0,001* | 3,36 ±<br>0,05 | <0,001* |
| 4 | 73,68<br>± 1,02 |  | 78,29<br>± 1,76 |  | 74,17<br>± 2,83 |  | 26,08 |  | 19,18<br>± 0,51 |  | 9,91 ±<br>0,49 |  | 0,99 ±<br>0,02 |  | 109,68<br>± 1,16 |  | 3,37 ±<br>0,01 |  |
| 5 | 67,48<br>± 2,45 |  | 76,52<br>± 0,81 |  | 69,99<br>± 1,50 |  | 8,50 |  | 18,37<br>± 0,36 |  | 9,53 ±<br>0,24 |  | 0,94 ±<br>0,04 |  | 101,06<br>± 0,9 |  | 3,28 ±<br>0,04 |  |
| 6 | 76,25<br>± 1,11 |  | 80,05<br>± 0,87 |  | 76,75<br>± 2,38 |  | 26,92 |  | 19,58<br>± 0,40 |  | 10,29<br>± 0,94 |  | 1,02 ±<br>0,02 |  | 124,01<br>± 0,96 |  | 3,4 ±<br>0,01 |  |

\* Statistically significant results; the ANOVA test was used for all variables except AST, for which the Kruskal-Wallis test was used.

**Table 6.** Serum biochemical parameters of female rats.

| Groups | Glucose (mg/dL) |  | Total cholesterol (mg/dL) |  | Triglycerides (mg/dL) |  | AST (U/L) |  | ALT (U/L) |  | Urea (mg/dL) |  | Creatinine (mg/dl) |  | Sodium (mEq/L) |  | Potassium (mEq/L) |  |
| --- | --- | --- | --- | --- | --- | --- | --- | --- | --- | --- | --- | --- | --- | --- | --- | --- | --- | --- |
|  | Mean<br>± SD | *p* | Mean<br>± SD | *p* | Mean<br>± SD | *p* | Mean<br>± SD | *p* | Mean<br>± SD | *p* | Mean<br>± SD | *p* | Mean<br>Rank | *p* | Mean ±<br>SD | *p* | Mean<br>± SD | *p* |
| 1 | 65,05<br>± 4,28 |  | 71,59<br>± 1,84 |  | 67,28<br>± 4,51 |  | 22,65<br>± 0,4 |  | 18,2 ±<br>0,67 |  | 18,2 ±<br>0,67 |  | 5,83 |  | 131,39<br>± 2,62 |  | 3,25 ±<br>0,06 |  |
| 2 | 65,74<br>± 2,6 |  | 76,39<br>± 3,96 |  | 70,23<br>± 1,83 |  | 36,07<br>± 1,04 |  | 18,69<br>± 0,9 |  | 18,69<br>± 0,9 |  | 13,25 |  | 123,47<br>± 1,86 |  | 3,28 ±<br>0,04 |  |
| 3 | 71,93<br>± 1,04 | <0,001* | 77,52<br>± 2,69 | <0,001* | 70,84<br>± 3,85 | 0,008* | 37,22<br>± 0,87 | <0,001* | 18,85<br>± 0,57 | 0,018* | 18,85<br>± 0,57 | 0,571 | 21 | <0,001* | 116,14 ±<br>3,71 | <0,001* | 3,34 ±<br>0,02 | <0,001* |
| 4 | 72,13<br>± 1,81 |  | 77,77<br>± 2,43 |  | 73,42<br>± 3,87 |  | 38,11<br>± 1,33 |  | 19,1 ±<br>0,59 |  | 19,1 ±<br>0,59 |  | 28 |  | 111,85 ±<br>1,04 |  | 3,35 ±<br>0,03 |  |
| 5 | 67,34<br>± 2,21 |  | 75,38<br>± 2,33 |  | 69,37<br>± 1,74 |  | 23,3 ±<br>0,43 |  | 18,13<br>± 0,5 |  | 18,13<br>± 0,5 |  | 11,67 |  | 103,71<br>± 0,91 |  | 3,28 ±<br>0,03 |  |
| 6 | 73,48<br>± 2,47 |  | 79,54<br>± 1,79 |  | 74,54<br>± 0,09 |  | 38,6 ±<br>1,23 |  | 19,26<br>± 0,4 |  | 19,26<br>± 0,4 |  | 31,25 |  | 126,35<br>± 1,06 |  | 3,37 ±<br>0,03 |  |

\*Statistically significant results; ANOVA test for all variables except creatinine using the Kruskal-Wallis test

**Table 7.** Post-hoc analysis of serum biochemical parameters of rats.

| Groups | *p* – value |  |  |  |  |  |  |  |  |  |  |  |  |  |  |  |
| --- | --- | --- | --- | --- | --- | --- | --- | --- | --- | --- | --- | --- | --- | --- | --- | --- |
|  | Male |  |  |  |  |  |  |  | Female |  |  |  |  |  |  |  |
|  | Glucose<sup>α</sup> | Total cholesterol<sup>α</sup> | Triglycerides<sup>β</sup> | AST<sup>δ</sup> | ALT<sup>β</sup> | Creatinine<sup>β</sup> | Sodium<sup>α</sup> | Potassium<sup>α</sup> | Glucose<sup>α</sup> | Total cholesterol<sup>β</sup> | Triglycerides<sup>β</sup> | AST<sup>β</sup> | ALT<sup>β</sup> | Creatinine<sup>δ</sup> | Sodium<sup>α</sup> | Potassium<sup>α</sup> |
| 1 – 2 | 0,99 | 0,55 | 0,091 | 0,003* | 0,002* | 0,09 | 0,009* | 0,97 | 0,82 | 0,003* | 0,13 | 0,00* | 0,19 | 0,052 | 0,00* | 0,87 |
| 1 – 3 | 0,13 | 0,17 | 0,33 | 0,003* | 0,001* | 0,00* | 0,00* | 0,17 | 0,65 | 0,00* | 0,072 | 0,00* | 0,08 | 0,008* | 0,00* | 0,07 |
| 1 – 4 | 0,00* | 0,52 | 0,00* | 0,004* | 0,00* | 0,00* | 0,00* | 0,00* | 0,56 | 0,00* | 0,003* | 0,00* | 0,02* | 0,004* | 0,00* | 0,06 |
| 1 – 5 | 0,96 | 0,66 | 0,16 | 0,05 | 0,22 | 0,54 | 0,00* | 0,94 | 0,84 | 0,01* | 0,283 | 0,25 | 0,82 | 0,059 | 0,00* | 0,13 |
| 1 – 6 | 0,00* | 0,01* | 0,00* | 0,004* | 0,00* | 0,00* | 0,095 | 0,00* | 0,025* | 0,00* | 0,001* | 0,00* | 0,01* | 0,004* | 0,00* | 0,03* |
| 2 – 3 | 0,10 | 1 | 0,63 | 0,42 | 0,75 | 0,032* | 0,012* | 0,12 | 0,052 | 0,46 | 0,75 | 0,046* | 0,66 | 0,066 | 0,00* | 0,048* |
| 2 – 4 | 0,00* | 1 | 0,005* | 0,14 | 0,35 | 0,01* | 0,00* | 0,00* | 0,051 | 0,36 | 0,11 | 0,00* | 0,27 | 0,006* | 0,00* | 0,048* |
| 2 – 5 | 0,99 | 1 | 0,77 | 0,003* | 0,036* | 0,27 | 0,00* | 0,99 | 1 | 0,51 | 0,66 | 0,00* | 0,12 | 0,677 | 0,00* | 1 |
| 2 – 6 | 0,00* | 0,77 | 0,00* | 0,143 | 0,017* | 0,00* | 0,047* | 0,00* | 0,021* | 0,04* | 0,032* | 0,00* | 0,128 | 0,004* | 0,017* | 0,016* |
| 3 – 4 | 0,99 | 1 | 0,017* | 0,62 | 0,54 | 0,61 | 0,166 | 0,99 | 1 | 0,86 | 0,19 | 0,117 | 0,50 | 0,07 | 0,001* | 1 |
| 3 – 5 | 0,89 | 0,78 | 0,44 | 0,003* | 0,01* | 0,002* | 0,001* | 0,10 | 0,19 | 0,16 | 0,45 | 0,00* | 0,53 | 0,022* | 0,00* | 0,02* |
| 3 – 6 | 0,43 | 0,70 | 0,00* | 0,62 | 0,036* | 0,092 | 0,003* | 0,46 | 0,72 | 0,25 | 0,064 | 0,018* | 0,27 | 0,012* | 0,00* | 0,088 |
| 4 – 5 | 0,00* | 0,93 | 0,003* | 0,004* | 0,00* | 0,001* | 0,00* | 0,01* | 0,021* | 0,12 | 0,043* | 0,00* | 0,01* | 0,004* | 0,00* | 0,024* |
| 4 – 6 | 0,18 | 0,93 | 0,051 | 0,68 | 0,13 | 0,23 | 0,00* | 0,009* | 0,87 | 0,25 | 0,57 | 0,38 | 0,66 | 0,254 | 0,00* | 0,88 |
| 5 – 6 | 0,00* | 0,10 | 0,00* | 0,004* | 0,00* | 0,00* | 0,00* | 0,002* | 0,01* | 0,01* | 0,01* | 0,00* | 0,00* | 0,004* | 0,00* | 0,01* |

\*Statistically significant results; <sup>a</sup>Games-Howel test; <sup>β</sup>LSD test; <sup>δ</sup>Mann-Whitney test.

The blood glucose levels of rats increased linearly with increasing doses of the test preparation in both male and female rats. Significant differences in blood glucose levels were found in male rats between group 1 compared to groups 4 and 6; group 2 compared to groups 4 and 6; and group 5 compared to groups 4 and 6. In female rats, significant differences in blood glucose levels were observed between groups 1, 2, and 5 compared with group 6, and between group 4 and group 5. The highest blood glucose level in the male rat group was 76.25 ± 1.11 mg/dL, while in the female rat group it was 73.48 ± 2.47 mg/dL. These values are still within the normal range for blood glucose in male rats (62.4-201.8 mg/dL) and in female rats (56.1-197.2 mg/dL(13).

The total cholesterol levels of the rats increased linearly with increasing doses of the test preparation, in both male and female rats. Significant differences in cholesterol levels were observed in male rats between groups 1 and 6. In female rats, significant differences were observed between group 1 and groups 2-6, and between group 6 and groups 2 and 5. The highest total cholesterol level in the male rat group was 80.05 ± 0.87 mg/dL, while in the female rat group it was 79.54 ± 1.79 mg/dL. These values are still within the normal range for total cholesterol in male rats (14.4-81.7 mg/dL) and in female rats (20.4-87.6 mg/dL(13).

Similar to total cholesterol levels, triglyceride levels in male and female rats also increased in direct proportion to the increase in the dose of the test preparation. In male rats, significant differences were found between group 1 and groups 4 and 6; between group 2 and groups 4 and 6; between group 3 and groups 4 and 6; and between group 5 and groups 4 and 6. In female rats, significant differences were found between group 1 and groups 4 and 6; between group 2 and group 6; and between group 5 and groups 4 and 6. The highest triglyceride level in the male rat group was 76.75 ± 2.38 mg/dL, while in the female rat group it was 74.54 ± 0.09 mg/dL. These values are still within the normal range for rat triglycerides, which is 30-409 mg/dL(14).

Liver function, assessed by AST and ALT levels, increased directly with the dose of the test preparation. In male rats, a significant difference in AST levels was found between group 1 and group 5 compared to all treatment groups. Similarly, significant differences in ALT levels were observed between group 1 and group 5 compared with all treatment groups, and between group 3 and group 6. The highest AST level was 38.76 ± 0.29 U/L, while the highest ALT level was 19.58 ± 0.4 U/L, found in group 6. In female rats, there was also a linear increase in AST and ALT levels with increasing doses of the test preparation. Significant differences in AST levels were found between group 1 and group 5 compared with all treatment groups; between group 2 and groups 3 and 4; between group 4 and group 5; and between group 6 and groups 2, 3, and 5. Meanwhile, a significant increase in ALT levels was observed in group 1 compared to groups 4 and 6, and in group 5 compared to groups 4 and 6. The highest AST and ALT levels were found in group 6, with values of 38.6 ± 1.23 U/L and 19.26 ± 0.4 U/L, respectively. Overall, the AST and ALT levels of all rats were within the normal range of 37-205 U/L and 6-114 U/L(14).

In the kidney function test, a significant difference between groups was observed only for creatinine. Creatinine levels increased in male and female rats in direct proportion to the increase in the dose of the test preparation. Significant increases were found in male rats between group 1 compared to groups 3, 4, and 6; group 2 compared to groups 4 and 6; group 3 compared to groups 5 and 6; and group 6 compared to groups 4 and 5. The highest creatinine level was found in group 6 at 1.02 ± 0.02 mg/dL (male rats) and 1 ± 0.01 mg/dL (female rats). The creatinine levels of all rats were within the normal range of 0.2-1.2 mg/dL(13).

In this study, serum electrolyte levels of Sodium (Na) and Potassium (K) were examined, which play an important role in maintaining body fluid balance, nerve and muscle function, and blood pressure regulation. The Na level in male and female rats decreased significantly, with the decrease inversely proportional to the dose of the test preparation. In male rats, significant differences were found between group 1 and groups 2-5; between group 2 and groups 4, 5, and 6; between group 5 and groups 3 and 4; and between group 6 and groups 3, 4, and 5. However, Na levels in group 6 did not differ significantly from those in group 1. In female rats, a significant decrease in Na levels was found in all comparisons between groups. The highest Na levels were found in group 1 male and female rats, at 128.74 ± 3.25 mEq/L and 131.39 ± 2.62 mEq/L, respectively. The average Na level for all rats was below the normal range of 139-155 mEq/L(14)

The K value in this study increased linearly with increasing dose of the test preparation. In male mice, significant increases were found in group 1 compared to groups 4 and 6; group 2 compared to groups 4 and 6; group 4 compared to groups 5 and 6; and group 5 compared to group 6. In female mice, significant differences were found between group 1 and group 6; between group 2 and groups 3, 4, and 6; and between group 5 and groups 3, 4, and 6. The highest K level was 3.4 ± 0.01 in male rats and 3.37 ± 0.03 mEq/L in female rats. The average potassium level for all mice was below the normal range of 3.6-7.7 mEq/L(14).

### Effects of NONB on the histology of liver, kidney, heart, lung, and spleen

In this study, histological observations were made on several vital organs (liver, kidney, heart, lung, and spleen) as shown in Figures 2–6. Bivariate analysis of histologic scores showed significant differences between groups in the liver, kidney, and spleen of male rats (Table 8). In female rats, significant differences were observed in the liver and spleen (Table 9).

**Figure 2.**
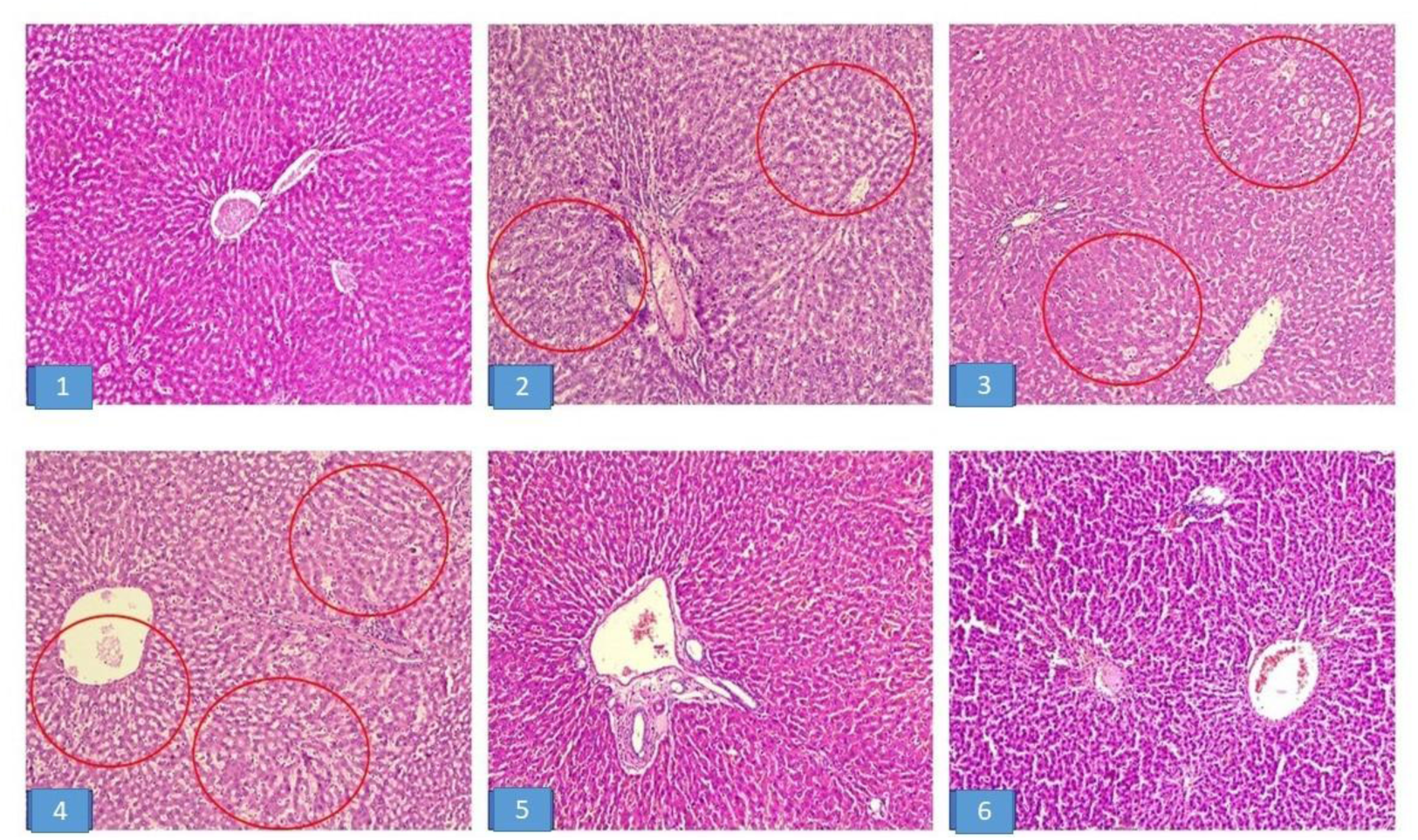
Normal liver histology (score 1) of rats in groups 1, 5, and 6; liver histology with multiple areas of fatty degeneration (red circle) in groups 2, 3, and 4 (score 3). (HE, 100x).

**Figure 3.**
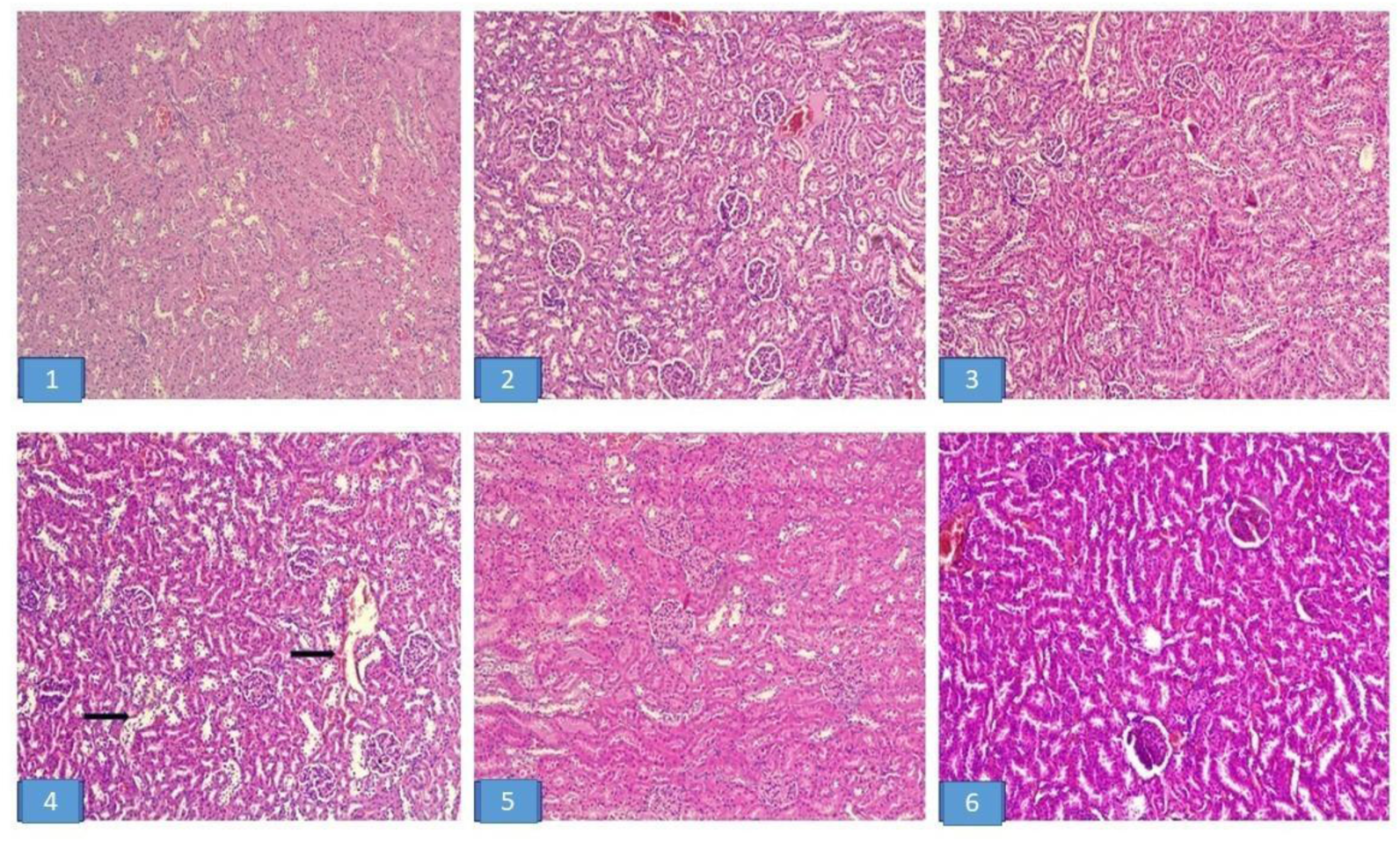
Normal histology of the kidney (score 1) of rats in groups 1, 2, 3, 5, and 6; proximal tubular dilation with epithelial degeneration (black arrow/score 2) in group 4. (HE, 100x).

**Figure 4.**
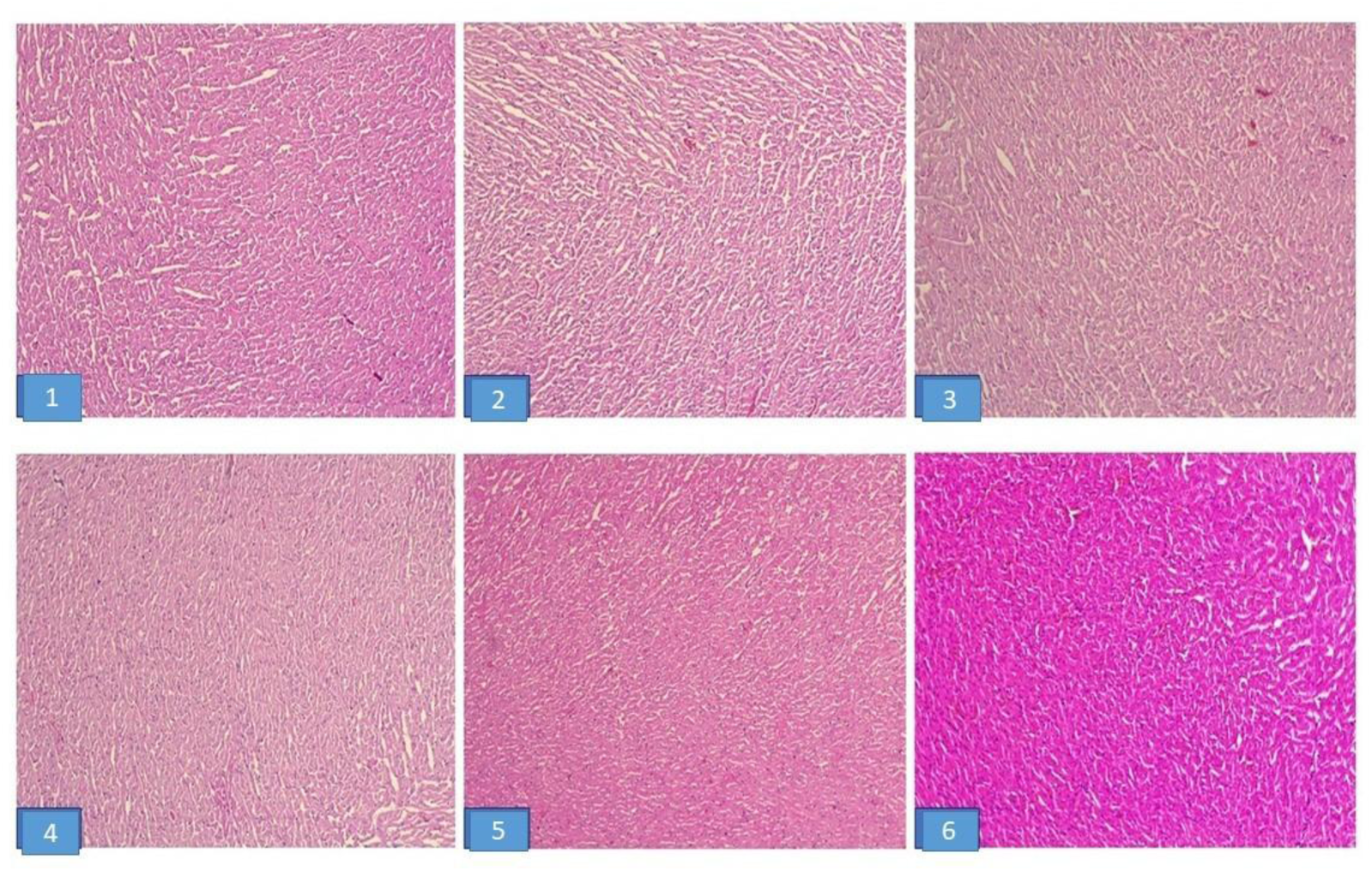
Normal histology of the heart (score 1) of rats in all groups. (HE, 100x)

**Figure 5.**
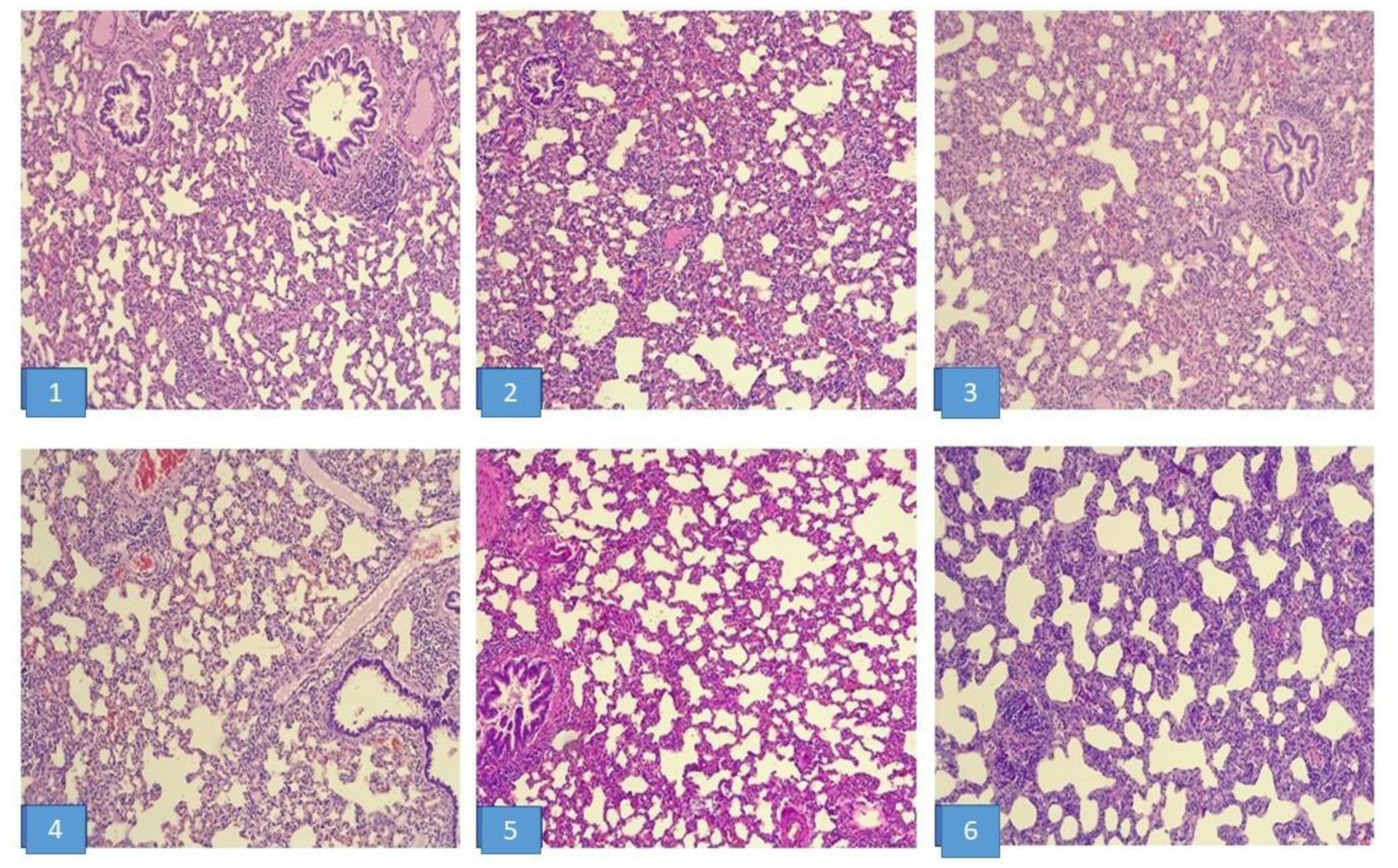
Normal histology of the lung (score 1) of rats in all groups. (HE, 100x)

**Figure 6.**
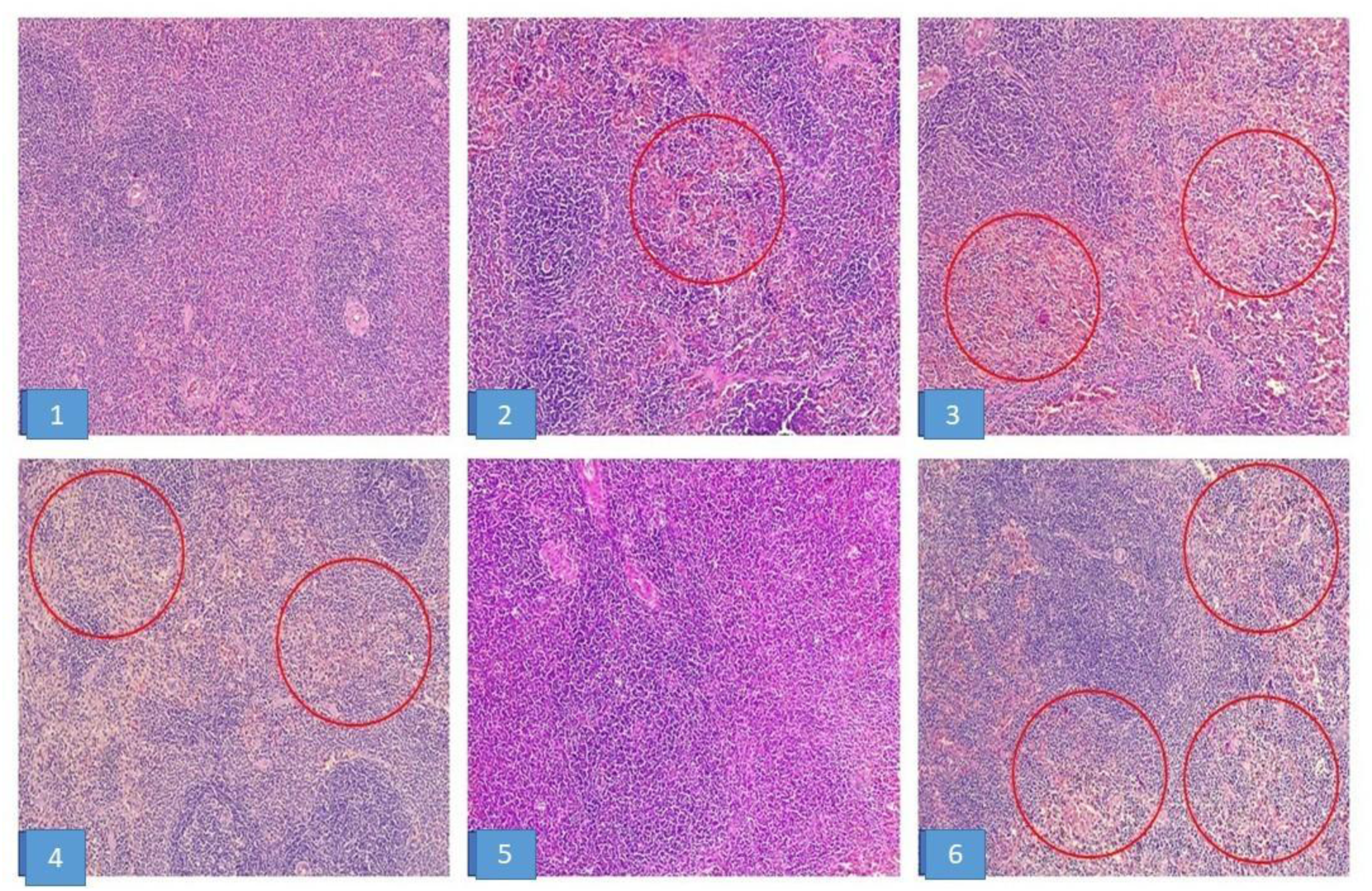
Normal spleen histology (score 1) of rats in groups 1 and 5; focal hemorrhage found in group 2; multifocal hemorrhage (red circle/ score 3) with hemosiderophage infiltration found in groups 3, 4, and 6. (HE, 100x).

**Table 8.**
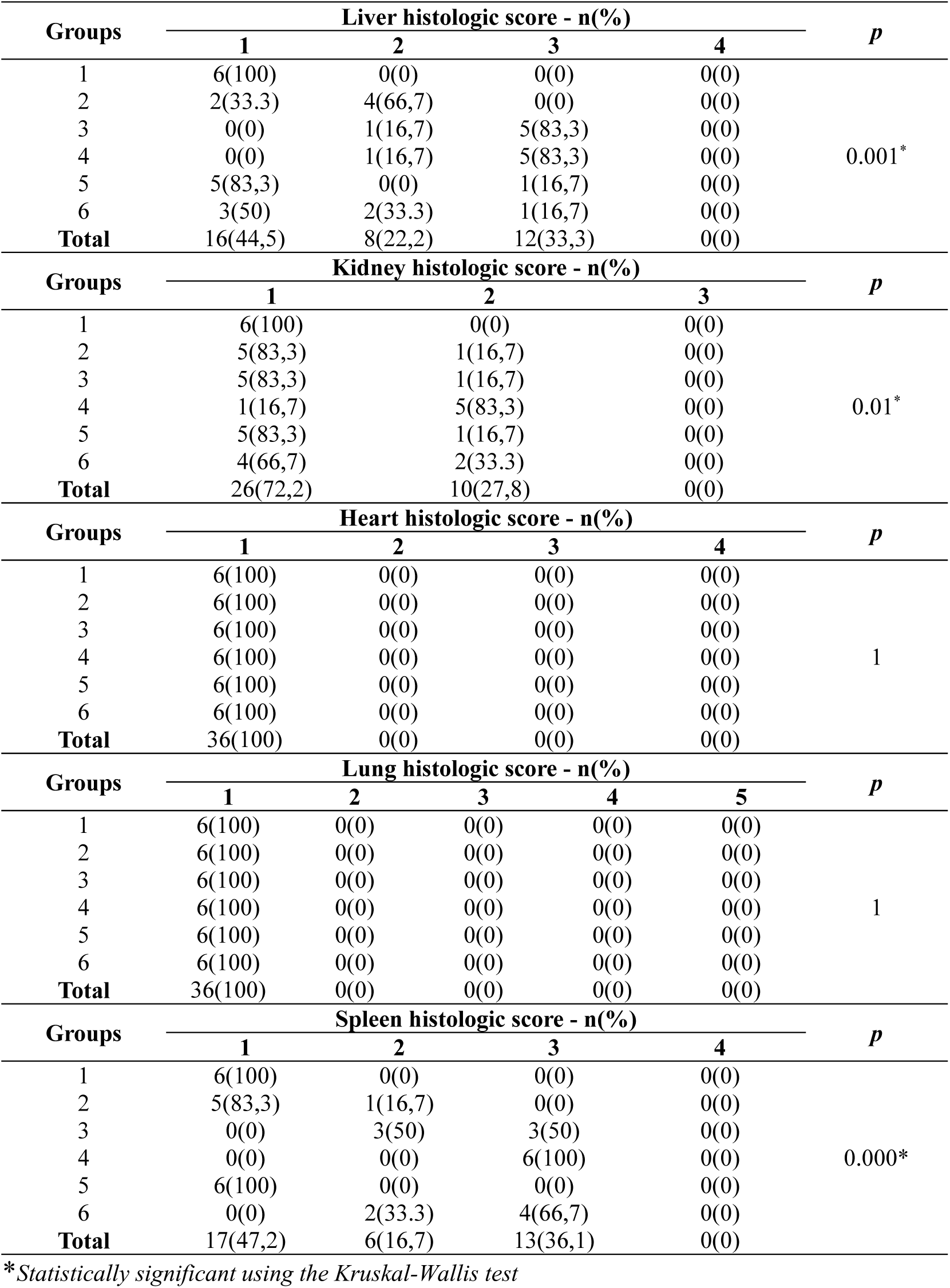
Histologic score of organs of male rats.

**Table 9.** Histologic score of organs of female rats.

| Groups | Liver histologic score - n(%) |  |  |  | *p* |  |
| --- | --- | --- | --- | --- | --- | --- |
|  | 1 | 2 | 3 | 4 |  |  |
| 1 | 6(100) | 0(0) | 0(0) | 0(0) | 0.005* |  |
| 2 | 4(66,6) | 1(16,7) | 1(16,7) | 0(0) |  |  |
| 3 | 1(16,7) | 1(16,7) | 4(66,67) | 0(0) |  |  |
| 4 | 0(0) | 6(100) | 0(0) | 0(0) |  |  |
| 5 | 4(66,7) | 2(33.3) | 0(0) | 0(0) |  |  |
| 6 | 3(50) | 1(16,7) | 2(33.3) | 0(0) |  |  |
| Total | 18(50) | 11(30,6) | 7(19,4) | 0(0) |  |  |
| Groups | Kidney histologic score - n(%) |  |  | *p* |  |  |
|  | 1 | 2 | 3 |  |  |  |
| 1 | 6(100) | 0(0) | 0(0) | 0.458 |  |  |
| 2 | 6(100) | 0(0) | 0(0) |  |  |  |
| 3 | 5(83,3) | 1(16,7) | 0(0) |  |  |  |
| 4 | 4(66,7) | 2(33.3) | 0(0) |  |  |  |
| 5 | 5(83,3) | 1(16,7) | 0(0) |  |  |  |
| 6 | 4(66,7) | 2(33.3) | 0(0) |  |  |  |
| Total | 30(83,3) | 6(16,7) | 0(0) |  |  |  |
| Groups | Heart histologic score - n(%) |  |  |  | *p* |  |
|  | 1 | 2 | 3 | 4 |  |  |
| 1 | 6(100) | 0(0) | 0(0) | 0(0) | 1 |  |
| 2 | 6(100) | 0(0) | 0(0) | 0(0) |  |  |
| 3 | 6(100) | 0(0) | 0(0) | 0(0) |  |  |
| 4 | 6(100) | 0(0) | 0(0) | 0(0) |  |  |
| 5 | 6(100) | 0(0) | 0(0) | 0(0) |  |  |
| 6 | 6(100) | 0(0) | 0(0) | 0(0) |  |  |
| Total | 36(100) | 0(0) | 0(0) | 0(0) |  |  |
| Groups | Lung histologic score - n(%) |  |  |  |  | *p* |
|  | 1 | 2 | 3 | 4 | 5 |  |
| 1 | 6(100) | 0(0) | 0(0) | 0(0) | 0(0) | 1 |
| 2 | 6(100) | 0(0) | 0(0) | 0(0) | 0(0) |  |
| 3 | 6(100) | 0(0) | 0(0) | 0(0) | 0(0) |  |
| 4 | 6(100) | 0(0) | 0(0) | 0(0) | 0(0) |  |
| 5 | 6(100) | 0(0) | 0(0) | 0(0) | 0(0) |  |
| 6 | 6(100) | 0(0) | 0(0) | 0(0) | 0(0) |  |
| Total | 36(100) | 0(0) | 0(0) | 0(0) | 0(0) |  |
| Groups | Spleen histologic score - n(%) |  |  |  | *p* |  |
|  | 1 | 0(0) | 3 | 4 |  |  |
| 1 | 6(100) | 0(0) | 0(0) | 0(0) | 0.000* |  |
| 2 | 5(83,3) | 1(16,7) | 0(0) | 0(0) |  |  |
| 3 | 0(0) | 3(50) | 3(50) | 0(0) |  |  |
| 4 | 0(0) | 0(0) | 6(100) | 0(0) |  |  |
| 5 | 6(100) | 0(0) | 0(0) | 0(0) |  |  |
| 6 | 0(0) | 2(33.3) | 4(66,7) | 0(0) |  |  |
| Total | 17(47,2) | 6(16,7) | 13(36,1) | 0(0) |  |  |

\*Statistically significant using the Kruskal-Wallis test

The results of the post-hoc analysis of histologic scoring with significant differences between groups are presented in Table 10. Significant differences in liver histology scores in male rats were found between group 1 compared to groups 2, 3, and 4; group 5 compared to groups 3 and 4; and group 4 compared to group 6. In female rats, significant differences in liver histology scores were observed between group 1 and groups 3 and 4, and between group 4 and groups 2, 5, and 6. Significant kidney histology scores were observed only in male rats in group 4, compared with groups 1, 2, 3, and 5. Significant spleen histology differences in male rats were found between group 1 compared to groups 2, 3, 4, and 6; group 2 compared to groups 4 and 5; and between group 5 compared to groups 3, 4, and 6. In female rats, significant differences in spleen histology scores were observed between group 1 and groups 2, 3, 4, and 6; between group 2 and groups 4 and 5; and between group 5 and groups 3, 4, and 6.

**Table 10.** Post-hoc analysis of the histologic score of organs in rats.

| Groups | *p - Value* |  |  |  |  |
| --- | --- | --- | --- | --- | --- |
|  | Male |  |  | Female |  |
|  | Liver | Kidney | Spleen | Liver | Spleen |
| 1 – 2 | 0.019* | 0.317 | 0.001* | 0.140 | 0.001* |
| 1 – 3 | 0.001* | 0.317 | 0.002* | 0.006* | 0.002* |
| 1 – 4 | 0.001* | 0.005* | 0.001* | 0.001* | 0.001* |
| 1 – 5 | 0.317 | 1.000 | 1.000 | 0.138 | 1.000 |
| 1 – 6 | 0.058 | 0.138 | 0.002* | 0.058 | 0.002* |
| 2 – 3 | 0.399 | 1.000 | 0.241 | 0.069 | 0.241 |
| 2 – 4 | 0.399 | 0.027* | 0.005* | 0.006* | 0.005* |
| 2 – 5 | 0.093 | 0.317 | 0.001* | 0.847 | 0.001* |
| 2 – 6 | 0.226 | 0.523 | 0.093 | 0.527 | 0.241 |
| 3 – 4 | 1.000 | 0.027* | 0.056 | 0.140 | 0.056 |
| 3 – 5 | 0.01* | 0.317 | 0.002* | 0.154 | 0.002* |
| 3 – 6 | 0.0198 | 0.523 | 0.575 | 0.221 | 1.000 |
| 4 – 5 | 0.01* | 0.005* | 0.001* | 0.019* | 0.001* |
| 4 – 6 | 0.019* | 0.093 | 0.138 | 0.021* | 0.056 |
| 5 – 6 | 0.338 | 0.138 | 0.002* | 0.715 | 0.002* |

\*Statistically significant using the Mann-Whitney test

## Discussion

Nitric Oxide (NO) is an endogenous free radical and a signaling molecule involved in various roles in the nervous, cardiovascular, reproductive, and immune systems(15). Because of its relative lipophilicity, neutral charge, and small size, NO can readily diffuse through the cell membrane without the need for receptors or channels(16). NO is known for its paracrine effects on the vascular wall and for its short biological half-life of 1-10 seconds, with a short diffusion distance of 50-1,000 µm(17).

Relevant studies have demonstrated the unique biological activities and chemical properties of specific gaseous signaling molecules in the human body, including NO, carbon monoxide, and hydrogen sulfide(18). Among the many gaseous molecular transmitters, NO exhibits highly prominent physiological and pathological effects in studies on various cells(19). Besides its important role in various physiological regulatory systems, many studies have confirmed the close link between NO and the incidence and pathogenesis of multiple diseases(20). Therefore, the use of NO for treatment has attracted widespread attention from scientists.

Researchers have attempted to develop NO donor delivery systems, including nano-delivery systems, which are expected to overcome NO’s short half-life (21), including those from IMI. IMI is developing NO nanobubble (NONB) technology that enables the encapsulation and controlled release of NO(22). This technology is expected to be an innovative solution for NO delivery, improving efficacy while minimizing side effects.

Based on the observations made, the NONB preparation administered at various doses did not cause the death of the experimental rats during the 90-day treatment period. The rat’s activity was also monitored and found to be normal, with no physical changes, signs, or symptoms of illness. Body weight increased proportionally with age, including the average weights of the liver, kidneys, heart, lungs, and spleen at the end of the study, and was also proportional to body weight. This condition indicates that the highest dose administered, 0.06 mL of NONB per day for 90 days, is still far below the lethal dose 50 (LD50).

The blood glucose levels of male and female rats in this study increased in direct proportion to the increase in the dose of NONB injected. This result is consistent with previous research showing that NO donors efficiently increase blood glucose levels that had been effectively decreased by *nitro-L-arginine methyl ester (L-NAME)*, an NOS inhibitor(23). On the other hand, previous studies have shown the opposite effect of NO on glucose metabolism. NO is the primary regulator of insulin secretion by the Langerhans islets, playing an important role in biphasic insulin secretion and exerting an insulin-stimulating effect(24). NO is involved in carbohydrate metabolism, and inhibition of the NO pathway leads to insulin resistance and Type 2 Diabetes Mellitus(25).

Controversy surrounding the role of NO in glucose metabolism may arise from its dual effects on glucose-stimulated insulin secretion. NO can act as both a regulator and a potential disruptor of glucose metabolism, depending on the concentration of exposure. Low to moderate concentrations can increase glucose uptake and utilization. In contrast, high concentrations can inhibit glucose oxidation, reduce insulin secretion, and decrease glucose uptake(24–26), as evidenced by elevated blood glucose levels. In this study, the increase in blood glucose levels in both male and female rats following NO injection did not return to baseline within 4 weeks of the last injection; instead, there was a slight, non-significant increase. These findings indicate that the effect of the highest NO injection dose (0.06 mL) persists even after exposure has been discontinued. However, in general, intravenous NO injection in this study was relatively safe for glucose metabolism, as blood glucose levels in all experimental rats remained within normal limits by the end of the study.

Total cholesterol and triglyceride levels in both male and female rats also increased in this study, in direct proportion to the increase in NONB dose. This condition differs from previous research findings that demonstrated the important role of NO in lipid metabolism regulation. NO has a beneficial effect on lipid metabolism by activating s*terol regulatory element binding protein (SREBP)-2*, and reduced NO bioavailability is associated with impaired lipid metabolism, increased levels of cholesterol, triglycerides, and LDL, and decreased levels of HDL(27–29). However, in vitro studies on AML-12 hepatocytes exposed to NO donors support these findings. Exposure of hepatocytes to NO donors increases lipid accumulation, depending on dose and duration. Donor NO causes an increase in the production of oxygen reactive species, particularly peroxides. At the same time, NO also increases GSH levels and activates antioxidant transcription factors such as *Hypoxia-inducible factor 1α (HIF1α)* and *Nuclear factor erythroid 2-related factor 2 (Nrf-2)*(30).

Similar to its effect on glucose metabolism, NO also has a dual effect on lipid metabolism. NO can act as a potential inhibitor of lipid peroxidation by scavenging peroxyl radicals and inhibiting peroxidase enzymes. However, NO can also be a potent oxidant through its reaction with superoxide to form peroxynitrite, a strong oxidant that can promote lipid peroxidation and oxidize lipid-soluble antioxidants(31). This condition is consistent with the findings of our study, which showed increased total cholesterol and triglyceride levels followed by fatty degeneration in rat hepatocytes. However, overall, the intravenous administration of the NONB test preparation up to the highest dose (0.06 mL) was still relatively safe, as the total cholesterol and triglyceride levels of all experimental rats remained within the normal range throughout the study.

AST and ALT levels are commonly used to evaluate liver function and detect liver damage or disorders. In this study, an increase in AST and ALT levels in rats was observed with increasing doses of NONB administered. These findings indicate a liver disturbance, confirmed by fatty degeneration of hepatocytes in the group that received NONB injections. The results of this study are consistent with previous research, both *in vitro* on AML-12 hepatocytes exposed to the NO donor *Diethylenetriamine-NONOate (DETA-NO)*(30) and *in vivo* in experimental rats exposed to sodium nitroprusside (SNP), an antihypertensive drug that induces NO production(32).

However, other studies have shown that NO has a protective effect on the liver. Research on CCl4-induced rats has demonstrated the hepatoprotective effect of NO donor *isosorbide-5-mononitrate*(33). Another study using NO V-PYRRO/NO donors also showed beneficial hepatoprotective effects on liver anatomy and physiology(34). The controversy surrounding existing research findings proves the complexity of NO’s role in hepatotoxicity. The source of NO production, the dose and duration of exposure, and micro environmental conditions are factors that influence NO’s role in liver condition and function. In this study, overall intravenous NONB exposure remained relatively safe, as increases in AST and ALT levels remained within the normal range, and liver histopathology improved in the satellite group after 4 weeks post-last injection.

In this study, kidney function was evaluated by examining urea and creatinine levels. No significant difference in urea levels was observed between the control group and the NONB-injected group. However, creatinine levels increased significantly in direct proportion to the increase in NONB dosage. The creatinine levels in the rats in this study remained within the normal range, indicating that the highest NONB dose (0.06 mL) did not cause kidney dysfunction. This condition was confirmed by kidney histology in male rats, which showed only mild damage. In contrast, no significant difference in kidney histology scores was observed between the control and NONB-injected groups in female rats.

The results of this study are consistent with previous research. NO is a vasodilator that plays an important role in renal hemodynamic homeostasis in both normotensive and hypertensive conditions(35). Research in rats has shown that both exogenous and endogenous NO exposure protect against ischemia/reperfusion-induced kidney tissue damage and dysfunction by suppressing endothelin-1 overproduction in post-ischemic kidneys(36). Additionally, clinical studies in patients undergoing coronary angiography (CAG) and percutaneous coronary intervention (PCI) have demonstrated that supplemental NO donor therapy reduces the incidence of contrast-induced nephropathy and decreases serum creatinine levels, regardless of administration orally or intravenously(37).

NO plays an important role in regulating sodium and potassium balance. NO affects transport, reabsorption, and both their effects on blood vessels and overall cardiovascular function(38,39). In this study, serum sodium levels decreased in the NONB-injected group. This condition may occur due to NO’s role in regulating renal blood pressure and sodium excretion, as well as hormonal regulation by aldosterone, vasopressin, angiotensin II, and endothelin(40). Donor NO, such as SNP, has been shown to increase renal sodium and water excretion, thereby decreasing serum sodium levels(41). In this study, the decrease in serum sodium levels was found to be temporary, as serum sodium levels in the satellite group returned to normal. Improved histological findings in the kidney also support this condition.

Unlike sodium, serum potassium levels in this study increased with increasing doses of NONB injection. These results are consistent with previous in vitro and in vivo studies demonstrating that NO donors activate KATP channels. This activation causes increased potassium efflux, which contributes to elevated serum potassium levels(42,43). Although serum sodium levels decreased and potassium levels increased, by the end of the study, both levels tended to approach the lower standard limit. These results prove that intravenous NONB injections generally do not affect electrolyte balance. This condition is also evident from the relatively normal histology of the kidneys, heart, and lungs.

In this study, pathological conditions were found in the spleens of rats injected with NONB i.v. linearly with increasing doses of the test preparation. Spleen histology shows bleeding with an infiltration of *hemosiderin-laden macrophages*. Hemosiderin is a form of iron storage, primarily derived from the breakdown of erythrocytes, which typically occurs in the red pulp of the spleen. Under normal conditions, hemosiderin can be found in the spleen in varying amounts depending on the species. Excessive amounts of splenic hemosiderin can be found in conditions of reduced erythropoiesis or rapid erythrocyte destruction in hemolytic anemia. Excessive hemosiderin in the spleen can also occur in chronic heart failure, iron dextran injections, focal accumulation of hematomas, infarcts, and bleeding due to trauma(44).

The increased amount of splenic hemosiderin in this study is likely due to focal accumulation of hematoma or bleeding resulting from hemodynamic instability induced by NONB injection. Excessive NO can increase intrasplenic microvascular pressure by relaxing arteries more than intrasplenic veins. This condition causes extravasation of intrasplenic fluid into the lymphatic system, which can contribute to splenic bleeding(45,46). In addition, overproduction of NO can lead to loss of lymphoproliferative response in the spleen, particularly in T lymphocytes, resulting in cell death through apoptosis or necrosis(47). In this study, no histopathologic evidence of cell death or necrosis was found in splenic tissue. However, further research is needed to determine the cause and mechanism of the multifocal hemorrhage and hemosiderosis observed.

The variables tested in this study showed that the condition of the satellite group was similar to that of the highest-dose treatment group (0.06 mL). These results indicate that the effect of intravenous NONB injection persists for 4 weeks after the last injection. This condition requires further research, including quantifying NO levels in the body, to determine whether the effect is due to still-high NO levels or irreversible tissue damage resulting from its metabolic reactions. NO has a very short half-life, only a few seconds in the body and about 2 ms in the vascular lumen(48). From this research’s perspective, using nanobubble technology, NO’s presence in the body will persist longer, with a longer half-life.

Overall, the results of the intravenous NONB injection study in rats at graded doses of 0.01 mL, 0.04 mL, and 0.06 mL did not result in any deaths during the 90-day treatment period. Biochemical parameters for liver function, lipid profile, kidney function, and serum electrolyte levels remain within normal limits, with slight deviations in sodium and potassium levels. This condition is also supported by histological findings in the liver, kidney, and spleen, which show mild to moderate damage, while the heart and lungs are normal. Thus, it can be concluded that the intravenous administration of NONB up to a dose of 0.06 mL in this subchronic toxicity test is still categorized as safe. Further research, with modifications to the test formulation, is highly recommended through the clinical trial stage in humans.

## Acknowledgment

The Indonesian Molecular Innovation Foundation funded this research in collaboration with the Faculty of Medicine Universitas Jenderal Soedirman.

